# Cooperative behavior evokes inter-brain synchrony in the prefrontal and temporoparietal cortex: A systematic review and meta-analysis of fNIRS hyperscanning studies

**DOI:** 10.1101/2021.06.03.446922

**Authors:** Artur Czeszumski, Sophie Hsin-Yi Liang, Suzanne Dikker, Peter König, Chin-Pang Lee, Sander L. Koole, Brent Kelsen

## Abstract

Single-brain neuroimaging studies have shown that human cooperation is associated with neural activity in frontal and temporoparietal regions. However, it remains unclear whether single-brain studies are informative about cooperation in real life, where people interact dynamically. Such dynamic interactions have become the focus of inter-brain studies. An advantageous technique in this regard is functional near-infrared spectroscopy (fNIRS) because it is less susceptible to movement artifacts than more conventional techniques like EEG or fMRI. We conducted a systematic review and the first quantitative meta-analysis of fNIRS hyperscanning of cooperation, based on thirteen studies with 890 participants. Overall, the meta-analysis revealed evidence of statistically significant inter-brain synchrony while people were cooperating, with large overall effect sizes in both frontal and temporoparietal areas. All thirteen studies observed significant inter-brain synchrony in the prefrontal cortex (PFC), suggesting that this region is particularly relevant for cooperative behavior. The consistency in these findings is unlikely to be due to task-related activations, given that the relevant studies used diverse cooperation tasks. Together, the present findings support the importance of inter-brain synchronization of frontal and temporoparietal regions in interpersonal cooperation. Moreover, the present article highlights the usefulness of meta-analyses as a tool for discerning patterns in inter-brain dynamics.

## Introduction

Human beings cooperate on small scales, like friends or families, and on larger scales, like nation states (Handley & Mathew 2020; Jaeggi & Gurven 2013). Nevertheless, there are many cases where cooperation fails, from marital arguments to political conflicts, leading to suboptimal outcomes for individuals and society. To understand the complexities of cooperation and help people realize more of their cooperative potential, it is helpful to obtain a better scientific understanding of cooperation.

One key scientific question is how cooperation is implemented in the brain. Over the last three decades, a large literature has emerged on social neuroscience (Cacioppo et al., 2000; Todorov et al., 2011; Schurz et al., 2021). Much of this research to date has relied on a single-brain approach as the dominant paradigm in contemporary neuroscience. In a typical social neuroscience study, a participant views social stimuli on a computer screen while her or his neural activations are being recorded with EEG or fMRI. A number of neural systems have been implicated in social cognition more generally, including the mirror neuron system and the mentalizing system. The former purportedly consists of the inferior frontal gyrus (IFG), inferior frontal lobule (IFL), and superior temporal gyrus (STG). The latter involves the temporoparietal junction (TPJ), precuneus, and prefrontal cortex (PFC; Rizzolatti & Fabbri-Destro; 2008; van Overwalle & Baetens, 2009).

One limitation of traditional social neuroscience research is that participants are not directly engaged in social interaction. To overcome this problem, researchers have moved toward a truly social, second-person neuroscience approach (Redcay & Schilbach, 2019; Schilbach et al., 2013). In second-person neuroscience, neural processes are examined within the context of a real-time reciprocal social interaction. Preliminary evidence has confirmed the added value of the second-person neuroscience approach by showing that specific neural signatures are only observable during ‘true’ social interaction (Tognoli et al., 2007).

Recent developments in neuroimaging have enabled so-called ‘hyperscanning’, whereby the activity of two or more brains can be assessed simultaneously while people are interacting (Dumas et al., 2010, Czeszumski et al., 2020). The resulting inter-brain activity is usually characterized in terms of the synchronization of the functional activity of the interacting brains. Hyperscanning has used a variety of neural imaging procedures, including electroencephalography (EEG), magnetoencephalography (MEG), functional magnetic resonance imaging (fMRI), and functional near-infrared spectroscopy (fNIRS) (respectively: Goldstein et al., 2018; Hirata et al., 2014; Koike et al., 2016; Scholkmann et al., 2013). Each apparatus and method has different advantages and disadvantages for hyperscanning (Czeszumski et al., 2020; Ayrolles et al., 2021). Hyperscanning research paradigms vary from studying coordinated finger movements (Tognoli et al., 2007), to real-life situations like playing guitar in a duet (Sanger et al., 2012) or studying multiple brains of high-school students inside the classroom (Dikker et al., 2017).).

So far, hyperscanning studies have revealed that inter-brain synchrony plays a crucial role in joint attention, interpersonal communication and coordination, cooperation, and decision-making (review: Czeszumski et al., 2020). Many hyperscanning studies have used spoken language during interactions between participants (Kelsen et al., 2020; Pérez et al., 2017; Li et al., 2021), ranging from knowledge sharing, cooperation, turn-taking, and naturalistic situations. Of the latter studies, many reported the emergence of inter-brain synchrony during interpersonal communication based on cooperative interaction in frontal and temporoparietal regions.

While the field is still young (Czeszumski et al., 2020), we conducted a meta-analysis (Zlowodzki et al., 2007) of fNIRS hyperscanning studies focusing on cooperative behavior. The present review focused explicitly on fNIRS studies for a number of reasons. The method of fNIRS is one of the most commonly used neuroimaging techniques in hyperscanning studies of cooperation (Kelsen et al., 2020), which is relatively insensitive to motion artifacts and capable of capturing inter-brain synchrony over longer periods (from seconds to minutes).

For example, social communication enhanced inter-brain synchrony during a turn-taking game (Nozawa et al., 2016). These and related findings suggest that inter-brain synchrony in frontal regions is associated with successful knowledge sharing and cooperative behavior using spoken language. Studies have additionally reported higher inter-brain synchrony in temporoparietal regions during teacher-student interactions (Zheng et al., 2018; Liu et al., 2019), cooperation (Xue et al., 2018; Lu et al., 2019b), and naturalistic discussion (Jiang et al., 2015).

In sum, many hyperscanning studies have examined the inter-brain dynamics associated with cooperative behavior. The findings appear to show some convergence, with inter-brain synchrony seemingly emerging in frontal regions. However, without quantitative integration through meta-analysis, it is not possible to determine the degree to which hyperscanning studies of cooperation have converging results. This question is of substantive theoretical interest, given the diverse paradigms used in hyperscanning studies in this area. More specifically, the cooperation tasks used varied considerably across studies, ranging from singing together to jointly solving a puzzle. This means that these tasks, aside from their cooperative nature, are unlikely to evoke shared neural activations based on low-level operational features. Thus, finding a common neuroanatomical site for inter-brain synchrony in these studies would provide relatively strong evidence for a general-purpose neural substrate for cooperative behavior. Our work had two aims: (1) to review the relevant literature and (2) to assess consistency in findings of inter-brain synchrony in different brain regions related to cooperative behavior.

## Methods

### Search strategy and inclusion criteria

We searched MEDLINE and SCOPUS databases for fNIRS hyperscanning studies of cooperation in accordance with preferred reporting items for systematic reviews and meta-analysis guidelines (PRISMA, Moher et al., 2009). Following consultation with a librarian, two authors independently conducted searches in September 2021 using keywords: ((hyperscanning OR “social neuroscience” OR fnirs) AND (interbrain OR inter-brain OR interpersonal OR interneural OR inter-neural OR synchron* OR coupling OR alignment OR “functional connectivity”) AND (cooperat* OR collaborat*)). Inclusion criteria included: fNIRS hyperscanning; cooperation/collaboration (where participants interacted to achieve a specific outcome such as solve a problem or puzzle or accomplish a particular result, thereby excluding turn-taking activities such as sequential counting, ultimatum game, prisoner dilemma and word games). Additionally, we excluded studies that focused on comparisons between genders, different levels of cooperation and did not report comparisons between cooperation and other conditions (cooperation or independent) or baseline. Discrepancies relating to inclusion were resolved through mutual discussion.

**Figure 1.**
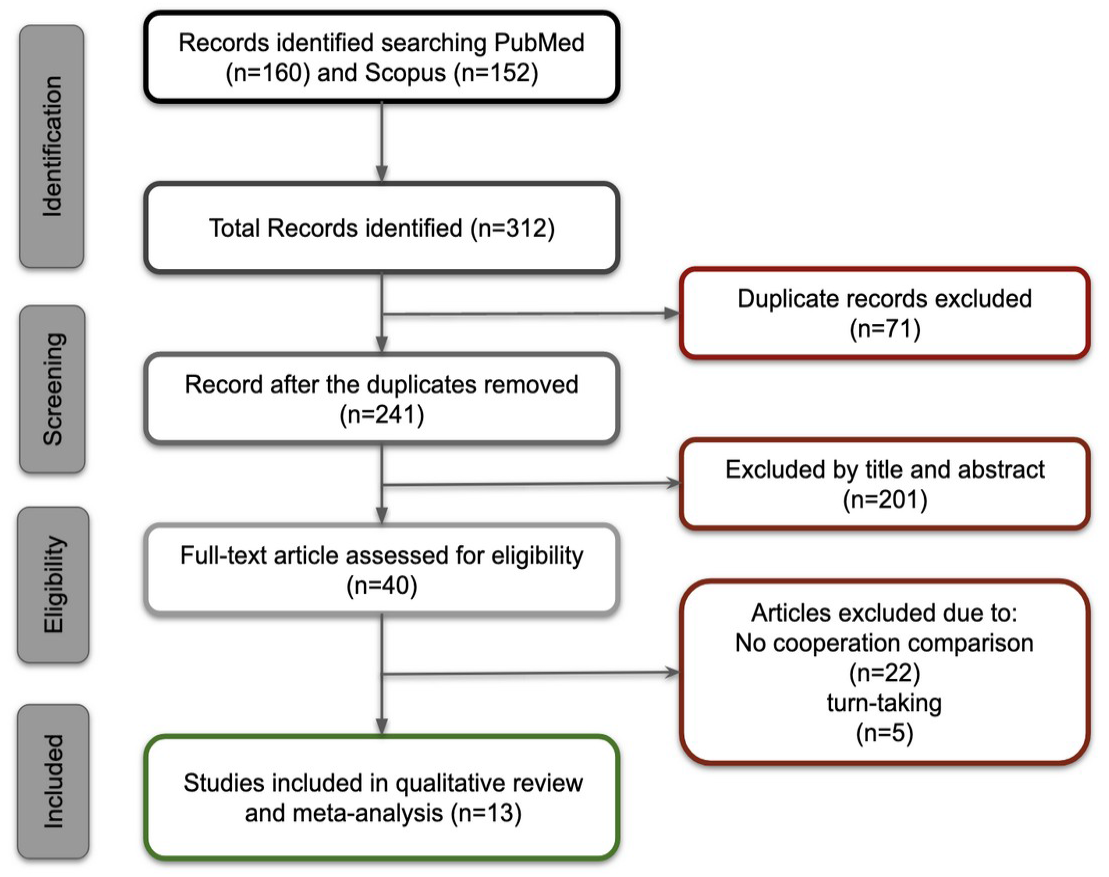
Flowchart of selection process.

### Statistical analyses

Because functional equivalence was not expected to hold across the included studies, and a common effect size could not be assumed, we performed a random-effects meta-analysis (Borenstein et al., 2009). We set the threshold for type I errors (alpha) at 0.05 and used effect sizes provided in the selected articles (if reported). We used the Psychometrica website (Lenhard and Lenhard, 2016) to estimate Cohen’s *d* from η^2^ (if available in the article), or we estimated Cohen’s *d* based on information provided in the article (statistical results)(Lipsey and Wilson, 2001). Further, we transformed effect sizes to Hedges’g; although similar to the classical Cohen’s *d*, it controls potential biases in studies with small sample sizes. If more than one comparison between cooperation and other conditions was present in the article, we chose the most orthogonal comparison. Furthermore, if more than one channel per region was reported, we selected the most central channel to the reported brain region. The heterogeneity across studies was gauged by Cochrane’s *Q, I*^*2*^, *τ*^*2*^statistics, and forest plots. We used Cochrane’s *Q* as a statistical test of the null-hypothesis of no heterogeneity, *I*^*2*^ to quantitatively estimate the variance between studies, and forest plots to visualize all effect sizes. In addition, we used funnel plots to assess publication bias. Publication bias concerns the elevated probability of studies reporting positive results being published. The tendency of journals to give preference to research showing positive findings means negative results may remain unpublished, leading to bias and an increased likelihood of false-positive outcomes (Zlodowski et al., 2007). Using Egger’s tests, we tested the funnel plot for symmetry and adjusted effect sizes with trim and fill analysis (Egger et al., 1997). Furthermore, we performed meta-regression analysis to test the influence of the variables Age, Gender and Language, Type of communication on overall effect sizes. All statistics were computed using the open-source JASP statistical computing environment (JASP Team, 2020).

## Results

We first present the results of the literature review and afterward the results of the meta-analysis of thirteen selected papers.

### Selected studies

The search resulted in selecting thirteen studies over the period 2016 to 2021, with an initial total of 888 participants and 847 once unusable data was removed (see Table 1). Nine studies were conducted in China, one in Japan and three were performed in the USA. Seven studies used verbal communication between acting participants during the investigation, while six studies did not. HbO measures were used due to increased sensitivity to blood flow, with pre-processing including low-pass filtering and global detrending. Eleven of the studies employed wavelet transform coherence (WTC; Grinsted et al., 2004) to convert the signal for inter-brain synchrony analysis, and two studies used correlation-based measures to estimate inter-brain synchrony.

**Table 1.**
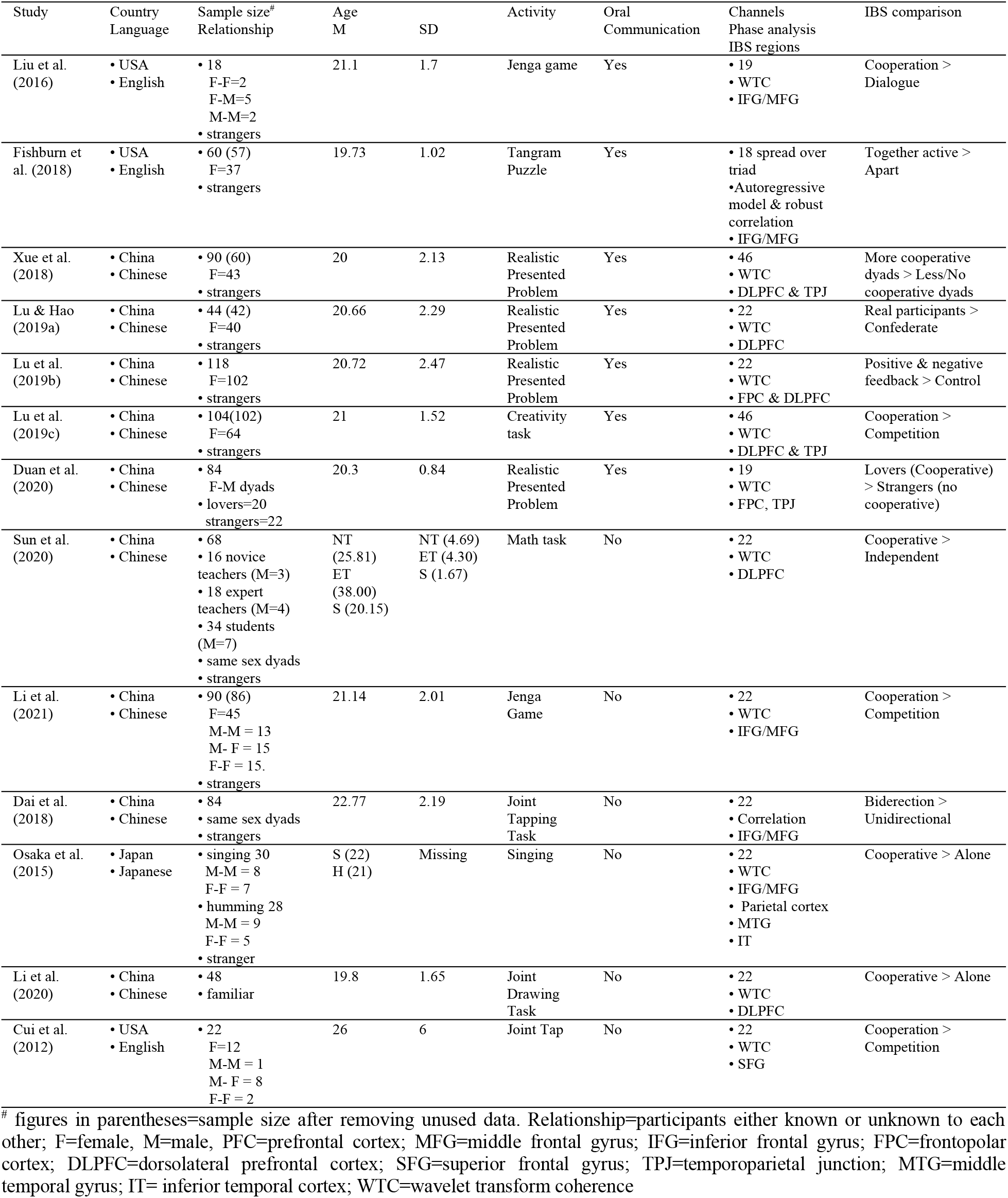
Selected studies

| Study | Country<br>Language | Sample size<sup>#</sup><br>Relationship | Age<br>M | SD | Activity | Oral<br>Communication | Channels<br>Phase analysis<br>IBS regions | IBS comparison |
| --- | --- | --- | --- | --- | --- | --- | --- | --- |
| Liu et al.<br>(2016) | • USA<br>• English | • 18<br>F-F=2<br>F-M=5<br>M-M=2<br>• strangers | 21.1 | 1.7 | Jenga game | Yes | • 19<br>• WTC<br>• IFG/MFG | Cooperation ><br>Dialogue |
| Fishburn et<br>al. (2018) | • USA<br>• English | • 60 (57)<br>F=37<br>• strangers | 19.73 | 1.02 | Tangram<br>Puzzle | Yes | • 18 spread over<br>triad<br>• Autoregressive<br>model & robust<br>correlation<br>• IFG/MFG | Together active ><br>Apart |
| Xue et al.<br>(2018) | • China<br>• Chinese | • 90 (60)<br>F=43<br>• strangers | 20 | 2.13 | Realistic<br>Presented<br>Problem | Yes | • 46<br>• WTC<br>• DLPFC & TPJ | More cooperative<br>dyads > Less/No<br>cooperative dyads |
| Lu & Hao<br>(2019a) | • China<br>• Chinese | • 44 (42)<br>F=40<br>• strangers | 20.66 | 2.29 | Realistic<br>Presented<br>Problem | Yes | • 22<br>• WTC<br>• DLPFC | Real participants ><br>Confederate |
| Lu et al.<br>(2019b) | • China<br>• Chinese | • 118<br>F=102<br>• strangers | 20.72 | 2.47 | Realistic<br>Presented<br>Problem | Yes | • 22<br>• WTC<br>• FPC & DLPFC | Positive & negative<br>feedback > Control |
| Lu et al.<br>(2019c) | • China<br>• Chinese | • 104(102)<br>F=64<br>• strangers | 21 | 1.52 | Creativity<br>task | Yes | • 46<br>• WTC<br>• DLPFC & TPJ | Cooperation ><br>Competition |
| Duan et al.<br>(2020) | • China<br>• Chinese | • 84<br>F-M dyads<br>• lovers=20<br>strangers=22 | 20.3 | 0.84 | Realistic<br>Presented<br>Problem | Yes | • 19<br>• WTC<br>• FPC, TPJ | Lovers (Cooperative)<br>> Strangers (no<br>cooperative) |
| Sun et al.<br>(2020) | • China<br>• Chinese | • 68<br>• 16 novice<br>teachers (M=3)<br>• 18 expert<br>teachers (M=4)<br>• 34 students<br>(M=7)<br>• same sex dyads<br>• strangers | NT<br>(25.81)<br>ET<br>(38.00)<br>S (20.15) | NT (4.69)<br>ET (4.30)<br>S (1.67) | Math task | No | • 22<br>• WTC<br>• DLPFC | Cooperative ><br>Independent |
| Li et al.<br>(2021) | • China<br>• Chinese | • 90 (86)<br>F=45<br>M-M = 13<br>M- F = 15<br>F-F = 15.<br>• strangers | 21.14 | 2.01 | Jenga<br>Game | No | • 22<br>• WTC<br>• IFG/MFG | Cooperation ><br>Competition |
| Dai et al.<br>(2018) | • China<br>• Chinese | • 84<br>• same sex dyads<br>• strangers | 22.77 | 2.19 | Joint<br>Tapping<br>Task | No | • 22<br>• Correlation<br>• IFG/MFG | Bidirection ><br>Unidirectional |
| Osaka et al.<br>(2015) | • Japan<br>• Japanese | • singing 30<br>M-M = 8<br>F-F = 7<br>• humming 28<br>M-M = 9<br>F-F = 5<br>• stranger | S (22)<br>H (21) | Missing | Singing | No | • 22<br>• WTC<br>• IFG/MFG<br>• Parietal cortex<br>• MTG<br>• IT | Cooperative > Alone |
| Li et al.<br>(2020) | • China<br>• Chinese | • 48<br>• familiar | 19.8 | 1.65 | Joint<br>Drawing<br>Task | No | • 22<br>• WTC<br>• DLPFC | Cooperative > Alone |
| Cui et al.<br>(2012) | • USA<br>• English | • 22<br>F=12<br>M-M = 1<br>M- F = 8<br>F-F = 2 | 26 | 6 | Joint Tap | No | • 22<br>• WTC<br>• SFG | Cooperation ><br>Competition |

<sup>#</sup> figures in parentheses=sample size after removing unused data. Relationship=participants either known or unknown to each other; F=female, M=male, PFC=prefrontal cortex; MFG=middle frontal gyrus; IFG=inferior frontal gyrus; FPC=frontopolar cortex; DLPFC=dorsolateral prefrontal cortex; SFG=superior frontal gyrus; TPJ=temporoparietal junction; MTG=middle temporal gyrus; IT= inferior temporal cortex; WTC=wavelet transform coherence

### Experimental designs

The conditions under which inter-brain synchrony occurred depended upon the experimental setup. Cooperative behavior is often studied with the use of games. Our search found three studies that used Jenga or Tangram puzzles to investigate inter-brain synchrony (Jenga: Liu et al., 2016; Li et al., 2021; Tangram: Fishburn et al., 2018). In the case of the Jenga game, these studies compared cooperative and competitive modes of building a tower, while solving a tangram puzzle was compared between together and apart conditions. On the one hand, multiple studies used different types of problem-solving tasks to study inter-brain synchrony. A set of studies (Lu et al., 2019a; Lu et al., 2019b; Duan et al., 2020; Xue et al., 2018) used realistically presented problem, where cooperation was facilitated by feedback and compared with situations where no feedback was provided. These studies utilized the presence of a third person (confederate) to create cooperative (feedback) and non-cooperative situations (no-feedback). This task closely resembles many everyday situations in which we solve problems together with the people surrounding us. They require communication and creativity; therefore, they are suitable for studying neural underpinnings of social interactions (inter-brain synchrony).

Lu et al., (2019c) used a creativity task in cooperative and competitive contexts. Participants in this study had to solve problems that required divergent thinking. Another aspect of cooperation was studied with a math problem task by Sun et al., (2020) by comparing cooperative with independent situations between a teacher and student (both adults). On the other hand, tasks that cooperatively require synchronization of behavior were selected. Two studies investigated synchronized taps between participants. In one of them, participants tried to synchronize their taps (cooperation) or be faster than the co-actor (competition) (Cui et al., 2012), while in the other study, bidirectional and unidirectional tapping was compared (Dai et al., 2018). Lastly, one study compared inter-brain synchrony in joint (synchronized) versus independent drawing (Li et al., 2020). In sum, various types of tasks were found to study cooperation and inter-brain synchrony with fNIRS. This suggests that many different cognitive functions were studied, and different brain regions were involved.

### Brain regions

The results of the studies we reviewed showed inter-brain synchrony in different parts of the brain. Studies reported parts of frontal and temporoparietal regions as sources of synchronization.

### Prefrontal Cortex (PFC)

All studies report different subregions of PFC to elicit more robust inter-brain synchrony in cooperative situations than the other conditions. Interestingly, different subparts of PFC were reported to be synchronized in different tasks. One set of studies (six: Xue et al., 2018; Lu et al., 2019a, 2019b, 2019c; Sun et al., 2020; Li et al., 2020) that required flexibility in solving a problem (realistic, creativity, and math problems) or drawing together show inter-brain synchrony in DLPFC. One of the primary functions of DLPFC reported in intra brain studies is cognitive flexibility related to attention switch (Monsell 2003).

Collaborative problem-solving tasks require focus switches between co-actors and the problem to solve, and inter-brain synchrony in DLPFC (dorso-lateral prefrontal cortext) may underpin these flexible attentional switches. Different subregions of PFC-IFG/MFG-show inter-brain synchrony during gamified tasks, like cooperative Jenga, tangram puzzle, and cooperative singing (four studies: Osaka et al., 2015, Liu et al., 2016, Fishburn et al., 2018, Li et al., 2021). These regions are involved in language processing, and inter-brain synchronization may facilitate cooperative behavior in tasks requiring a lot of verbal communication to solve (Jenga (with verbal communication) and Tangram puzzle; Liu et al., 2016, Fishburn et al., 2018). However, inter-brain synchronization in IFG/MFG (inferior frontal gyrus, middle frontal gyrus) was also reported in cooperative Jenga play without verbal communication (Li et al., 2021). Further research is needed to resolve the role of verbal communication in the Jenga task.One could compare cooperative Jenga play with and without verbal communication to gain more insight into the function of inter-brain synchrony in IFG/MFG.

Another subpart of PFC that shows inter-brain synchrony is SFG (superior frontal gyrus). We identified one experiment that showed higher inter-brain synchrony for cooperative joint tap when compared with competitive (Cui t al., 2012). Lastly, we found that FPC (frontopolar cortex) also shows inter-brain synchrony during cooperative realistic problem solving, suggesting that it is not only PFC that shows inter-brain synchrony. Taken together, we found that most of the studies show inter-brain synchrony in PFC, and that tasks requiring different cognitive functions elicit inter-brain synchrony in different subparts of PFC.

### Temporoparietal regions

Four of the included studies show inter-brain synchrony in temporoparietal regions. It is important to note that these four studies are not different studies from the studies discussed above, but they show inter-brain synchrony in temporoparietal regions in addition to PFC. Three out of four show inter-brain synchrony in the TPJ (temporoparietal junction) while participants solve realistic or creativity problems (Xue et al., 2018, Lu et al., 2019c, Duan et al., 2020). TPJ is involved in many different tasks that require the theory of mind (Schurz et al., 2014), which is essential for successful interpersonal interactions as cooperative problem solving (Rilling et al., 2004). Therefore, the results of selected studies extend past research by showing inter-brain synchrony in TPJ. Furthermore, these studies show inter-brain synchrony in both frontal and temporoparietal regions, suggesting the existence of a PFC-TPJ inter-brain network that facilitates cooperative behaviors. However, more evidence (studies) is required to test that interpretation. In addition to the PFC-TPJ connection, we identified one study that links PFC (IFG/MFG) with the temporal lobe (IT and MTG; inferior temporal cortex, middle temporal gyrus) during cooperative singing (Osaka et al., 2015).

Taken together, the selected studies pointed in the direction that inter-brain synchrony in prefrontal and temporoparietal regions plays a crucial role in cooperation. To test that further, we performed a meta-analysis of the selected studies.

**Figure 2.**
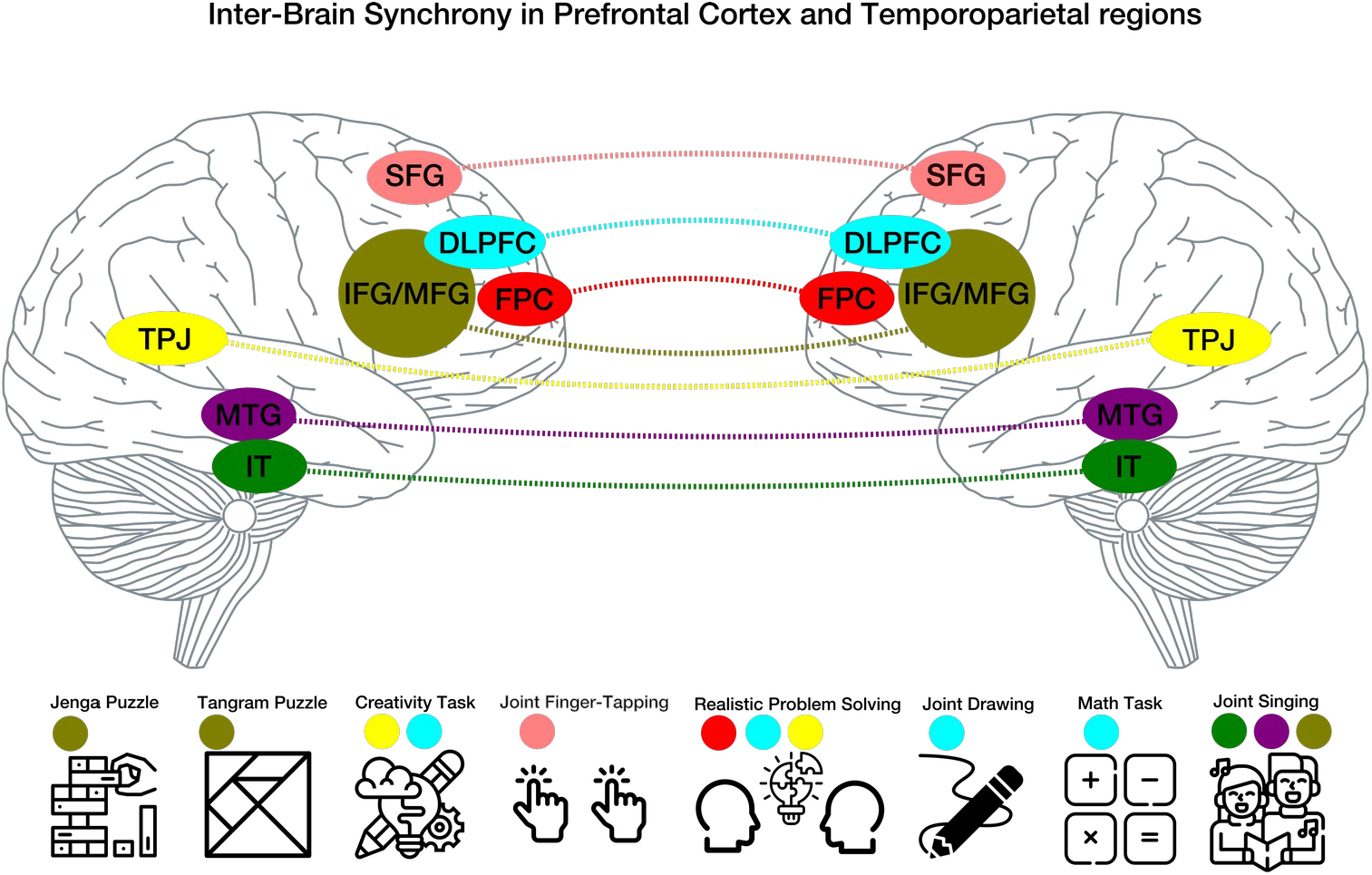
Inter-brain synchrony in different parts of the prefrontal and temporoparietal cortex in various tasks used to study cooperation.

#### Meta-analysis

A random-effects model for all twenty one experimental conditions across the thirteen studies reported a significantly large overall effect size (*g*=1.98, 95% CI [1.47, 2.49], n=21, *z*=7.68, *p*<0.001). Cochran’s Q-statistic (Q=469.72, *p*<0.001) showed significant variation around the weighted average effect for the studies included. The proportion of observed variance was significantly high at *I*^*2*^=98.6 (> 75 representing large heterogeneity), and a scaled measure of dispersion between true effect sizes of the studies was τ^2^=1.29 (Higgins & Thompson, 2002). These results suggest that the selected studies had an overall large effect size for comparison between cooperative and non-cooperative conditions. Furthermore, the variance between studies was high, suggesting that nearly all variance between studies was not due to chance. Visual inspection of the funnel plot and Egger’s test (z=7.22, p<0.001) indicated significant asymmetry. However, a follow-up trim and fill analysis resulted in the same effect size and confidence intervals (g=1.98, 95%CI [1.47, 2.49]).

We performed meta-regression examinations to test whether any independent variables (Age, Gender, Language, Type of Communication) affected our analysis. Wald tests demonstrated no significant association between observed inter-brain synchrony and independent variables overall. Chinese was used as the reference language. We found that, Age (*Beta*=0.12, *S*.*E*.=0.27, *z*=.43, *p*=0.66), Gender (*Beta*=-0.78,*S*.*E*.=2.45, *z*=-0.32, *p*=0.75) Communication (*Beta*=0.78, *S*.*E*.=1.4, *z*=.56, *p*=0.58) and Language (English) (*Beta*=0.12, *S*.*E*.=0.95, *z*=0.12, *p*=0.9), and (Japanese) (*Beta*=0.29, *S*.*E*.=0.92, *z*=0.3, *p*=0.77) all displayed insignificant results. The results of meta-regression analysis suggest that Age, Gender, Type of communication, and Language differences did not modulate overall effect sizes for the included studies.

**Figure 3.**
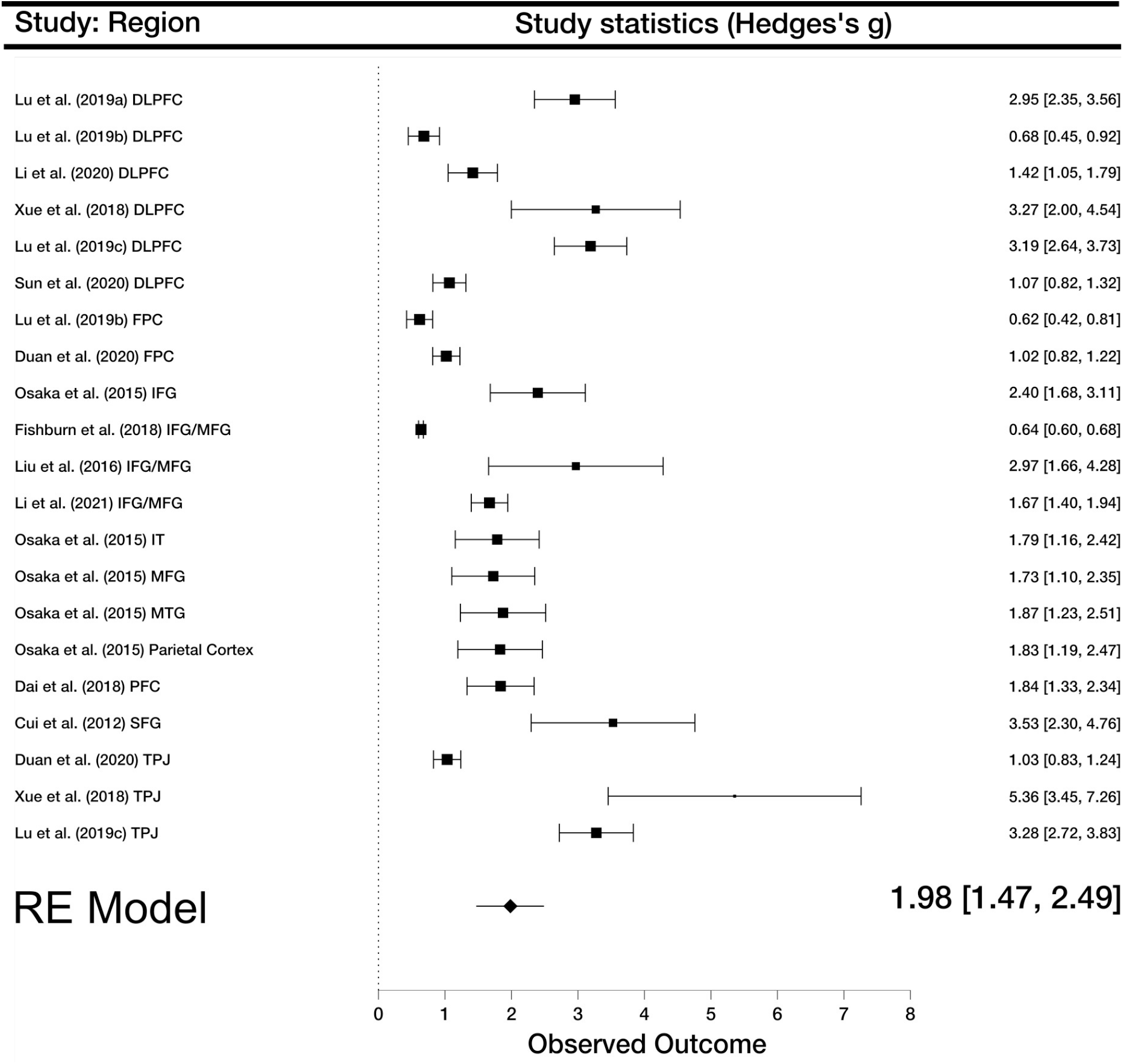
Forest plot of all included studies. Boxes represent effect sizes and whiskers confidence intervals.

## Discussion

When people cooperate, their neural activity will tend to become mutually synchronized. This inter-brain synchrony during cooperation tasks has become the focus of a growing number of hyperscanning studies. In the present article, we conducted a systematic review and meta-analysis of fNIRS hyperscanning studies of cooperation. We located thirteen relevant studies with a total of 890 participants. The results of our meta-analysis revealed significant overall effect sizes for inter-brain synchrony in both frontal and temporoparietal regions. All studies observed significant inter-brain synchrony in the prefrontal cortex (PFC). This consistency is remarkable, considering that the included studies used various cooperation tasks, such as realistic problem solving, joint drawing, and the Jenga puzzle. It thus appears that PFC has general relevance for cooperative behavior that cannot be reduced to task-specific elements.

The findings of the present meta-analysis are broadly consistent with the findings of previous single-brain studies implicating prefrontal regions in tasks requiring social interaction, coordination, and cooperation (Stallen & Sanfel, 2013). The present findings not only confirm these earlier findings from single-brain recordings but show that they are part of a broader pattern indicating that prefrontal regions are not just activated within individual brains operating separately from another. Instead, prefrontal regions are mutually activated in a synchronized fashion in the brains of interaction partners, becoming coupled in their functioning. Hyperscanning studies thus complement and extend traditional social neuroscience studies that were conducted within the single-brain paradigm.

The present work has limitations. First, the present meta-analysis included a relatively low number of studies. The studies had a relatively high number of participants, which affords better statistical power. Still, the limited number of studies makes it hard to estimate the effects of between-study characteristics. Second, the present meta-analysis was restricted to a single neuroimaging method, fNIRS, which has limited spatial resolution. In the same line, the placement of recording channels is not standardized; therefore, it is difficult to compare different studies. It hence remains essential to compare the present findings to other neuroimaging methods, like fMRI. Third, the meta-analysis revealed a high variance between studies that cannot be explained by chance. More work is needed to understand the sources of this variance, which is likely due to the large variety of conditions used in different studies. Fourth and last, the present meta-analysis may be contaminated by reporting bias, given that published studies tend to report only statistically significant comparisons of neural recordings. It is important to note that the last limitation is not a limitation per se of our work but a more general limitation of many neuroimaging studies that the field should address. We propose that no significant channels/comparisons should be reported in supplementary materials with all statistics values. It will allow for collecting more evidence and improve future meta-analyses. Additionally, this problem may be overcome in future work by creating better infrastructures for data sharing and open science practices (Pavlov et al., 2020).

## Conclusion

Human beings are a cooperative species. The present research uncovered some of the neural foundations of this human ability to cooperate by conducting the first systematic review and quantitative meta-analysis of fNIRS hyperscanning of cooperative behavior. The results showed that cooperation is consistently associated with inter-brain synchrony in frontal and temporoparietal areas, suggesting that inter-brain neural alignment in these regions underlies cooperative behavior in humans. These findings underscore the importance of meta-analyses in detecting patterns across studies and elucidating the neural basis of semi-naturalistic cooperative behavior.

